# Sleep deprivation constrains dynamic configurations of integrated and segregated brain states impacting cognitive performance

**DOI:** 10.1101/2025.10.21.683658

**Authors:** N. Cross, F. Borgetto, M. Uji, A. Jegou, A. Nguyen, K. Lee, U. Aydin, A. Perrault, C. Grova, T.T. Dang-Vu

**Affiliations:** Brain and Mind Centre, The School of Psychology, The University of Sydney, Australia; Department of Health, Kinesiology and Applied Physiology, Center for Studies in Behavioral Neurobiology, Sleep, Cognition, and Neuroimaging Lab, Concordia University, Montreal, Canada; Centre de Recherche de l’Institut Universitaire de Geriatrie de Montreal, CIUSSS Centre-Sud-de-l’Île-de-Montréal, Montreal, Canada; PERFORM Centre, Concordia School of Health, Concordia University, Montreal, Canada; RIKEN Center for Brain Science, Saitama, Japan; Department of Physics, Multimodal Functional Imaging Lab, Concordia University, Montreal Canada; Department of Psychiatry, Yale University School of Medicine, New Haven, USA; School of Psychology and Clinical Language Sciences, University of Reading, Reading, UK; Woolcock Institute of Medical Research, Macquarie University, Sydney, Australia; Department of Biomedical Engineering, Neurology and Neurosurgery, Multimodal Functional Imaging Lab, McGill University, Montreal, Canada

## Abstract

The breakdown of cognitive control following sleep deprivation is widely recognised, but the physiological mechanisms and brain signatures that produce this vulnerability have not been resolved. Effective cognition relies on large-scale brain networks flexibly reconfiguring between states of integration and segregation. Here we combined functional magnetic resonance imaging (fMRI), electroencephalography (EEG), and electrocardiography (ECG) collected during cognitive tasks under rested wakefulness, after sleep deprivation, and following a recovery nap to test the hypothesis that sleep deprivation constrains this dynamical repertoire and disrupts its physiological regulation. Using time-resolved functional connectivity and graph theory, we show that sleep deprivation increases the distribution of connections across networks, while reducing the temporal variability of between-network connectivity. Furthermore, dynamic fluctuations between integrated and segregated modes of network topology were dampened, with brain regions spending more time in intermediate configurations and showing greater instability of mode transitions. These alterations were tightly linked to behavioural impairment: participants who exhibited greater contraction toward intermediate topologies also showed poorer task accuracy and slower responses. Under well-rested conditions, thalamic activity peaked prior to transitions into integrated states and was suppressed during transitions into segregated states, consistent with a coordinating role in cortical dynamics. Sleep deprivation weakened and delayed this thalamic coupling. Finally, global and regional fMRI fluctuations were elevated after sleep loss, becoming decoupled from cardiac physiology while more strongly coupled to EEG delta power, further linking reduced arousal to constrained network flexibility. Together, these findings show that sleep deprivation narrows the brain’s dynamical repertoire, due to disrupted thalamic regulation and changes to the physiological integration with cortical networks.

## Introduction

The efficient execution of cognitive tasks is thought to rely on a dynamic balance between network segregation (reflected in the distinct activity of specialised cortical systems), and network integration, which supports the coordination of information flow across distributed regions. This integration– segregation balance has been proposed as a fundamental organizing principle of brain function, critical for flexible cognitive operations (Tononi et al. 1994; Sporns et al. 2004). Under conditions of reduced vigilance, such as prolonged wakefulness, endogenous fluctuations in cortical and subcortical activity become more pronounced (Wong et al. 2014; Cross et al. 2021). These fluctuations are associated with decreased network stability and may impair the brain’s ability to maintain effective integration across regions, potentially contributing to the cognitive control deficits observed following sleep loss.

Prior studies using functional magnetic resonance imaging (fMRI) have shown that sleep deprivation (SD) disrupts connectivity within and between large-scale brain networks (Yeo et al. 2015, Cross et al. 2021) and reduces global modularity in resting-state activity (Ben-Simon et al. 2017). However, relatively few studies have examined how SD alters brain connectivity during the performance of cognitively demanding tasks, particularly those requiring sustained cognitive control. In our previous work, we showed that brain signal fluctuations due to SD were associated with a cortex-wide increase in functional integration, primarily driven by elevated integration within cortical networks (Cross et al. 2021). This shift disrupted the typical balance between within- and between-network connectivity and was tightly linked to both impairments and recovery in cognitive control (Cross et al. 2021).

However, this previous work probed static functional connectivity patterns, averaged over the entire scanning time. Growing evidence indicates that large-scale brain networks reconfigure over shorter timescales from tens of seconds to a few minutes (Chang et al 2010, Handwerker 2012). Time-resolved analyses of fMRI data reveal that the brain transitions between transient states characterised by variable levels of connectivity strength and topological organisation (Lurie et al. 2019). These dynamic shifts are thought to reflect time-sensitive modulations in inter-regional communication, supporting ongoing cognitive processes (Fries et al. 2015). Specifically, the brain appears to cycle between relatively integrated modes, marked by strong global connectivity and network cohesion, and segregated modes, typified by increased modularity and local processing efficiency (Shine et al. 2016).

These transitions may be especially relevant in the context of sleep deprivation, where arousal-related endogenous fluctuations may perturb the frequency, duration, or structure of these dynamic modes. We previously found that heightened endogenous fluctuations in neural activity are closely linked to cognitive deficits and an altered balance of integrated and segregated brain activity (Cross et al. 2021, PLOS Biology). These cortical fluctuations occurred in the context of altered coordination by the thalamus, which plays a central role in sustaining arousal and constraining the engagement of distributed cortical assemblies (Edlow et al., 2012). Global BOLD fluctuations are tightly coupled to physiological rhythms, including slow EEG activity and autonomic signals such as heart rate variability and respiration (Tyvaert et al. 2008, Yuan et al. 2016), both of which are sensitive to sleep loss. Together, these observations highlight the thalamus and global physiological fluctuations as potential mechanisms through which sleep deprivation disrupts the balance of large-scale integration and segregation.

In the present study, we extend previous work by investigating dynamic functional connectivity during cognitive tasks performed under well-rested and sleep-deprived conditions. We aimed to characterise how SD influences the temporal expression and organisation of integrated and segregated brain modes and to determine whether alterations in these dynamic properties are linked to changes in cognitive performance. By capturing the moment-to-moment fluctuations in network topology, this approach may provide a more sensitive index of brain function under conditions of reduced arousal and help elucidate the mechanisms of information processing underlying cognitive vulnerability to sleep loss.

## Results

To assess the impact of sleep deprivation on dynamic brain connectivity during task performance, we analysed fMRI data from 20 participants who performed three cognitive tasks inside the MRI scanner under rested wakefulness, sleep deprivation, and following a recovery nap. EEG was recorded simultaneously to monitor vigilance and arousal. A sliding-window approach was applied to estimate time-resolved functional connectivity from fMRI data (task blocks regressed out), which was then characterised using graph theory measures of integration and segregation. Specifically, we calculated the participation coefficient (between networks, BT), which quantifies the distribution of a parcel’s connections between networks, and the within-module degree z-score (within networks, WT), which indexes the relative strength of within-network connectivity, for each of 400 cortical parcels (Schaeffer et al. 2018) across all time windows (Figure 1A).

**Figure 1.**
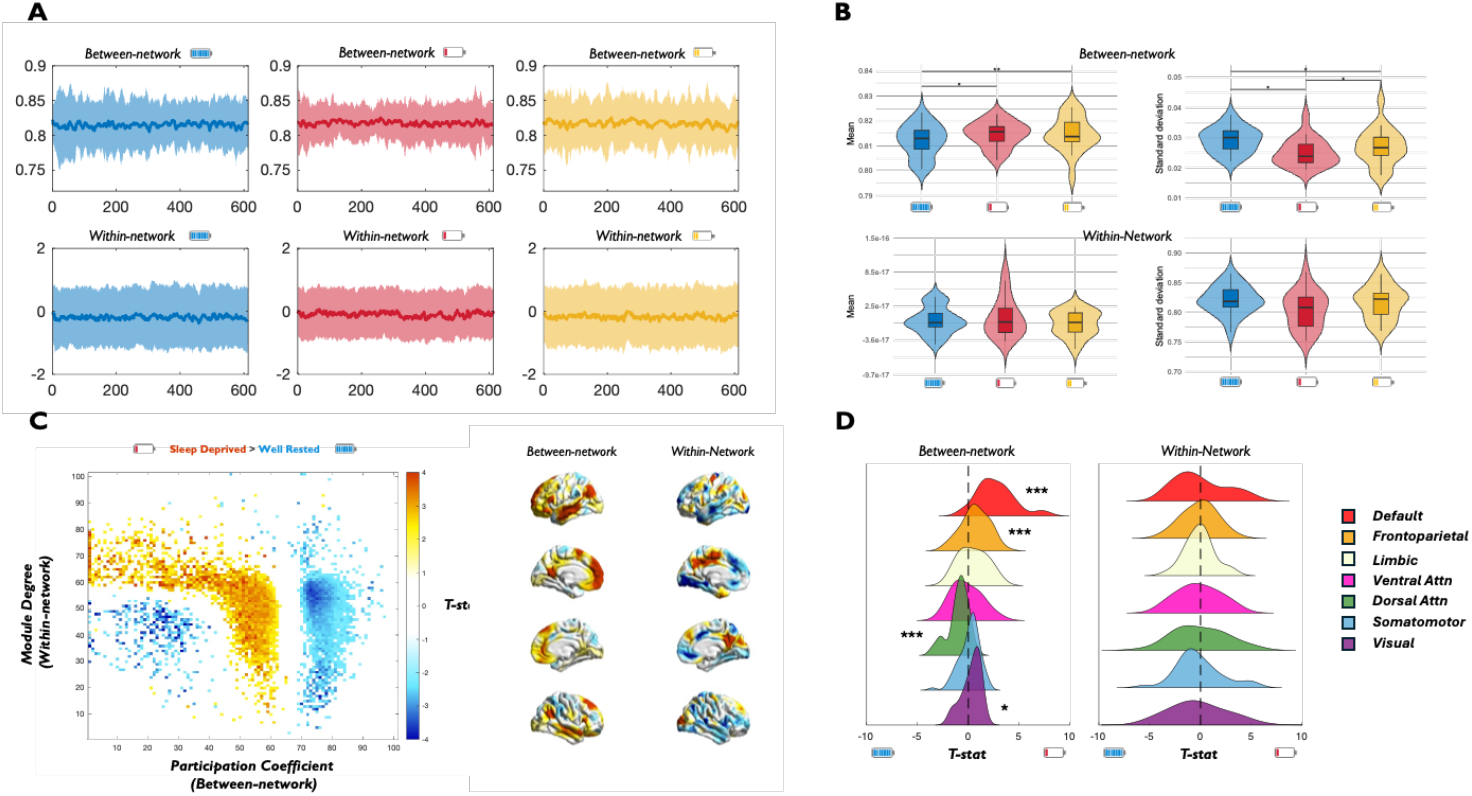
Sleep deprivation alters dynamic configurations of integration and segregation in the cerebral cortex. **A**| Exemplar timeseries of Between-network and Within-network connectivity for 1 participant across the Well-Rested, Sleep-Deprived and Post-Nap conditions. Solid Line represents mean of 400 parcels, shading ±1 sd. **B**| The group-level mean and standard deviation of Between-network and Within-network connectivity for 400 parcels across 613 timepoints (26min) in all participants. **C**| Regions of the 2D joint histogram (proportion of time spent in bins of BT and WT) that were significantly different between Sleep Deprived and Well Rested vigilance states during cognitive task performance blocks (paired-samples t-test). Coloured points indicate regions that survived false discovery correction (FDR, p < 0.05): red/yellow, increased frequency during Sleep Deprived state; blue/light blue, increased frequency during Well Rested state. Surface projections of cortical parcels associated with higher Between-network (left) or Within-network (right) connectivity during the Sleep-Deprived state, when compared the Well Rested state: default mode regions showed elevated between-network connections during sleep deprivation, whereas within-network connections were elevated in posterior cingulate and decreased in visual and medial temporal regions. **D**| Changes in Between-network connectivity within the 7-Yeo functional networks demonstrated significant increases in DMN, Frontoparietal and Visual networks, while a decrease in the Dorsal Attention network. No significant changes were observed in within network connectivity. *p<0.05, **p<0.01, ***p<0.001

Across the cortex, sleep deprivation was associated with a minor increase in the temporal mean BT (t_(20)_ = 2.2, p = 0.039), indicating more evenly distributed connections across networks (Figure 1B). This effect was sustained even following the recovery nap. By contrast, the temporal variability of BT was reduced: the standard deviation of BT across time windows was lower during sleep deprivation compared to both rested wakefulness (t_(20)_ = −5.4, p = 2.9 × 10^−5^) and the recovery nap (t_(20)_ = −2.0, p = 0.048), suggesting a dampening of dynamic fluctuations in between-network connectivity. In contrast to these between-network effects, there were no reliable differences in WT across conditions (Figure 1B), indicating that relative within-network connectivity remained stable.

To further characterise these dynamics, we applied the cartographic profiling method introduced by Shine et al. (2016) to construct a joint histogram of within- and between-network connectivity, representing the joint distribution of within- and between-network connectivity across all parcels and time windows (Figure 1C). This revealed a significant shift in the balance of topological states following sleep deprivation: brain parcels spent more time in intermediate regions of the cartographic space (neither high or low integration, Figure 1C) and in states reflecting high segregation (top left of the cartographic space). At the network level, the effects were more closely aligned with the observations in the temporal mean of BT. Notably, parcels within the default mode, frontoparietal, and visual networks spent more time with higher BT in the sleep-deprived state compared with rested wakefulness, while only the dorsal attention network exhibited decreases in BT (Figure 1D). No changes in WT were observed between the WR and SD conditions at the network level (Figure 1D). This indicates that while brain parcels spent more time in segregated regions of the cartographic space (Figure 1C), this was distributed evenly across all networks.

Following the recovery nap, we observed a partial reversal of the sleep deprivation pattern. The cartographic profile showed less time spent in intermediate topological states and a relative shift back toward more extreme integrated configurations (Supplementary Figure 1A). At the network level, changes in between-network connectivity were also evident: the default mode network (DMN) showed a significant decrease in participation coefficient, while all other Yeo-7 networks exhibited significant increases relative to the sleep-deprived state (Supplementary Figure 1B).

Overall, these complementary results demonstrate that sleep deprivation disrupts the normal balance of network topology by increasing average between-network connectivity while simultaneously constraining the temporal variability of these connections, leading to a collapse toward intermediate states (neither fully integrated nor segregated) that is only partially restored after recovery sleep.

### Fluctuations between integrated and segregated modes are reduced after sleep deprivation

Dynamic changes in network topology were further examined by quantifying time-resolved patterns of within- and between-network connectivity. For each time window, the cartographic profile was constructed by jointly mapping within-module degree (WT) and participation coefficient (BT) across all regions. To identify recurring modes of network organisation, all time-windowed cartographic profile sets were clustered using k-means (k = 2), yielding two distinct modes: an integrated mode, characterised by higher BT values, and a segregated mode, marked by lower BT and higher WT values (Figure 2A).

**Figure 2.**
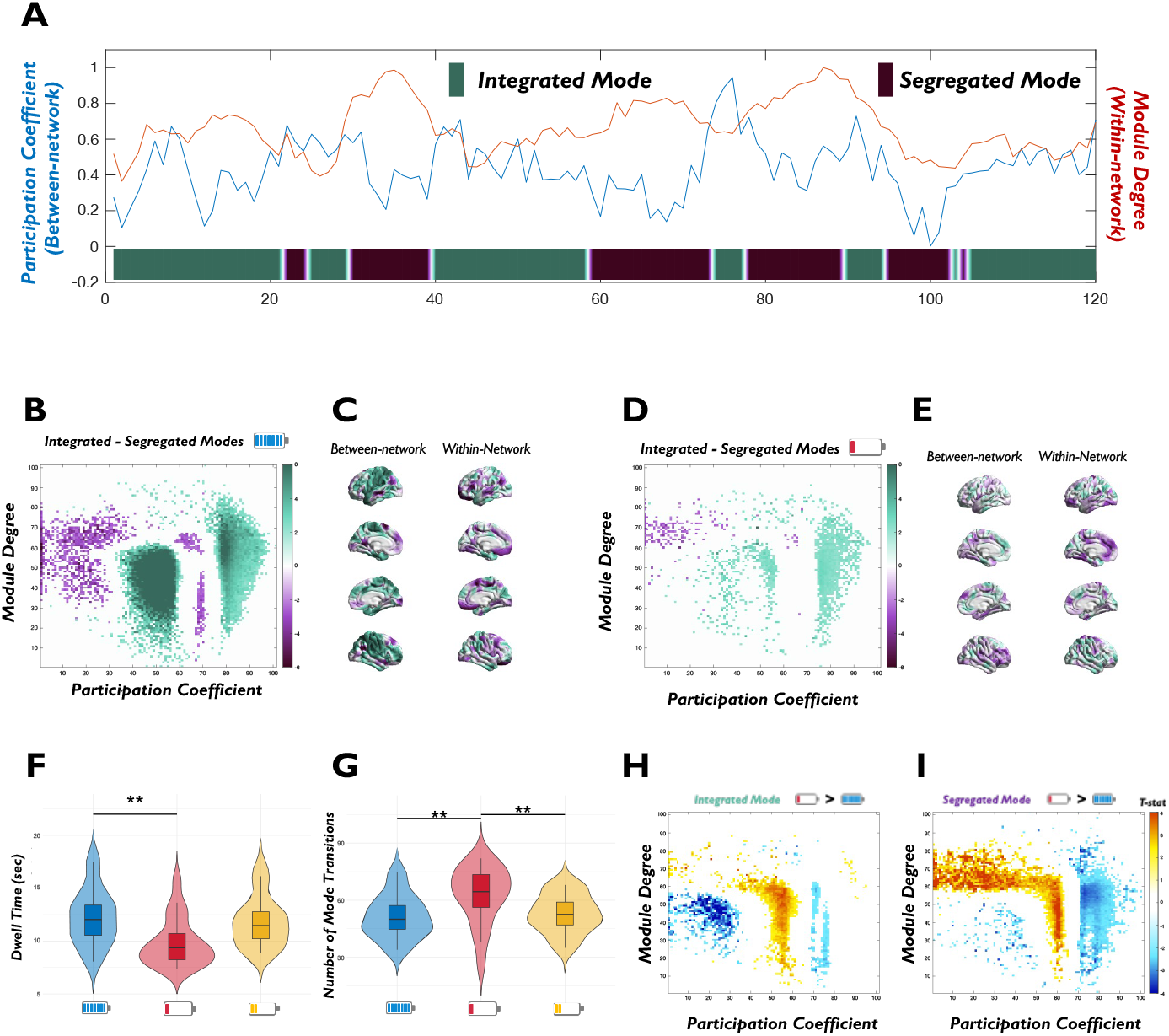
Fluctuations between integrated and segregated brain modes are dampened after sleep deprivation. **A**| Dynamic fluctuations in within- and between-network connectivity for a representative subject. Blue line reflects participation coefficient (BT), red line reflects within-module degree (WT), and overlaid state labels indicate transitions between integrated (green) and segregated (purple) brain modes determined by k-means clustering (k=2) of the cartographic profile. **B**| Group-level cartographic profiles, showing the differences in joint distribution of WT and BT values between integrated and segregated modes in the Well Rested (left) and Sleep Deprived conditions (right). Colour intensity indicates the frequency of regions occupying each topological state (Green = higher during integrated modes, Purple = higher during segregated modes, coloured points p_FDR_<0.05). **C**| Mean dwell time spent in each integrated and segregated modes and the number of transitions between modes across conditions represented. Bars show mean ± SEM across participants. **D**| Group-level cartographic profiles, showing the differences in joint distribution of WT and BT values between the Well Rested and Sleep Deprived conditions during integrated (left) and segregated (right) modes (Yellow = higher during Sleep Deprivation, Blue = higher during Well Rested condition, coloured points p_FDR_<0.05).

At the group level, cartographic profiles revealed clear differences between the two modes (Figure 2B). Integrated states were characterised by stronger BT and reduced WT, while segregated states showed very low BT and higher WT. Importantly however, the distribution of these states differed across conditions. In the Rested Wakefulness condition, the separation between integrated and segregated modes within the cartographic profile was more pronounced than after Sleep Deprivation (Figure 2B). Consistent with this reduced distinction, group-level analyses showed that sleep deprivation also increased the number of transitions between modes (t_(20)_ = 2.7, p = 0.013) and shortened the dwell time within each state (t_(20)_ = −3.0, p = 0.007), reflecting greater instability in the balance between integrated and segregated brain dynamics (Figure 2C).

Sleep deprivation dampened the extremes of both modes, reducing occupancy of highly integrated bins during the integrated mode and of highly segregated bins during the segregated mode—i.e., a contraction toward intermediate WT–BT values (Figure 2D). These results indicate that sleep deprivation disrupts the normal balance of large-scale brain dynamics, biasing the system toward a more persistent, segregated configuration at the expense of flexible switching between integrated and segregated network modes.

### Fluctuations in integration and segregation after sleep deprivation are associated with impaired cognition

Task performance was impaired following sleep deprivation, with accuracy reduced and response times slowed relative to rested wakefulness, and partial recovery observed after the nap (Figure 3A,C). To test whether these behavioural changes were linked to alterations in large-scale network dynamics, we compared changes in the cartographic profile with task performance. Across participants, reconfiguration of the joint WT–BT distribution between rested and sleep-deprived conditions was significantly correlated with changes in both accuracy and reaction time (Figure 3B,D), such that individuals whose brain parcels spent more time in intermediate regions of the cartographic space exhibited poorer task performance.

**Figure 3.**
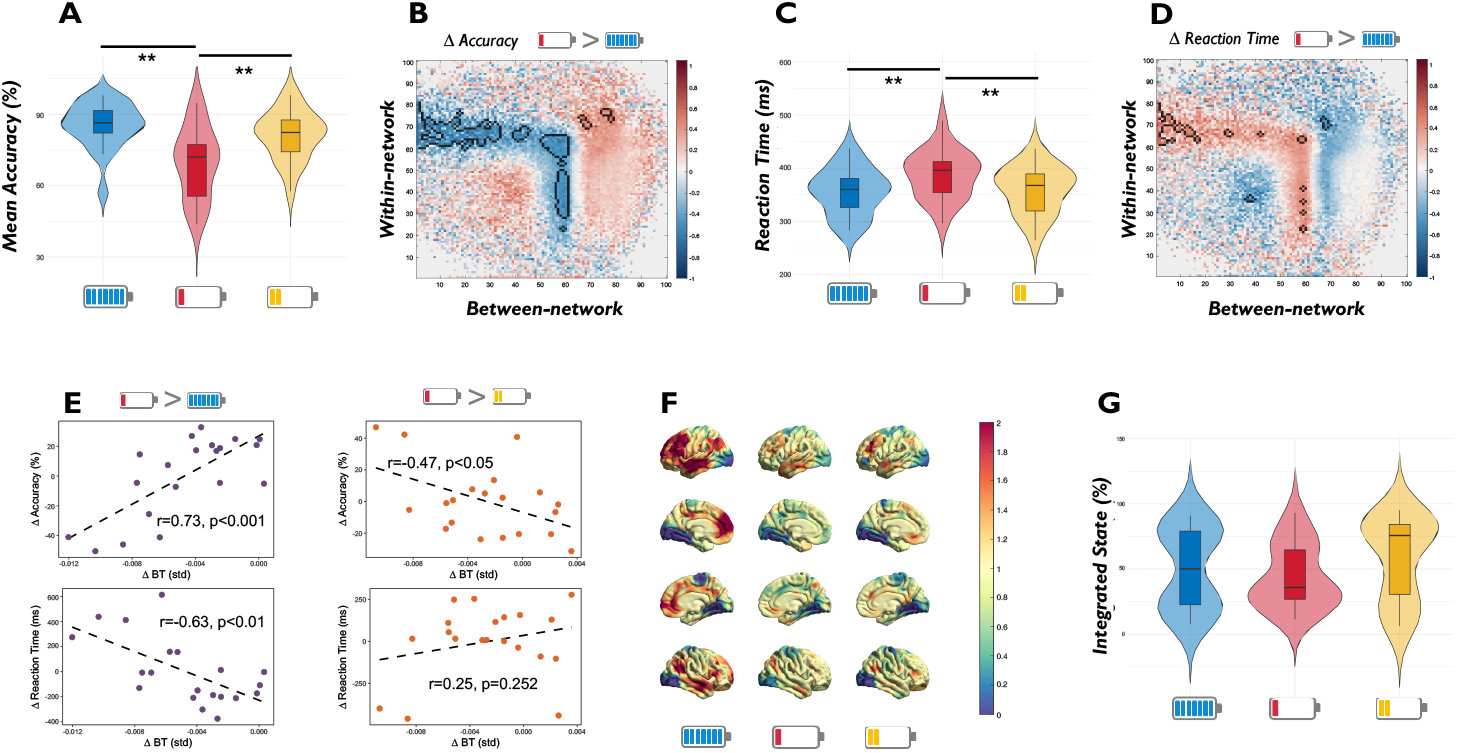
Fluctuations in configurations of integration and segregation after sleep deprivation are associated with impaired cognition. **A**| Task accuracy across Rested Wakefulness, Sleep Deprivation, and Post-nap recovery conditions. Violin plots show the distribution across participants, with black lines indicating the median. **B**| Differences in the cartographic profile between Sleep Deprivation and Rested Wakefulness correlated with changes in task accuracy. The heatmap shows correlation coefficients between topological reorganisation (WT– BT joint space) and behavioural scores, with warm colours indicating positive associations and cool colours negative associations. Outlined regions indicate clusters surviving significance thresholding (p < 0.05, FDR corrected). **C**| Task reaction time across conditions, plotted as in (A). **D**| Correlation between reconfiguration of brain network dynamics and changes in task reaction time, displayed as in (B). **E**| Association between temporal variability in between-network connectivity (ΔBTstd) and task performance. Scatterplots show correlations between changes in the standard deviation of participation coefficient (BT) and changes in accuracy and reaction time, for both WR–SD (left) and SD–PN (right) contrasts. **F**| General linear model (GLM) analysis showing regions where fluctuations in participation coefficient (BT) tracked the occurrence of behavioural lapses across tasks. Warmer colours indicate stronger positive associations, cooler colours indicate negative associations (unthresholded maps shown). **G**| Relationship between brain modes and lapses. Violin plots show the percentage of lapses and errors occurring during integrated (versus segregated) modes per condition, with black lines indicating the median across participants. Sleep deprivation increased the likelihood that lapses occurred during integrated modes and reduced the proportion during segregated modes, with partial recovery following the nap.

Variability in between-network connectivity across time was then evaluated in relation to behavioural performance. Reductions in the temporal standard deviation of the participation coefficient (ΔBTstd) were associated with lower accuracy (ρ = 0.73, p = 2.88 × 10^−4^) and slower responses (ρ = −0.63, p = 3.03 × 10^−3^) after sleep deprivation, while greater recovery of variability after the nap was linked to improved accuracy (ρ = 0.47, p = 0.035) but not reaction time (ρ = 0.25, p = 0.283, Figure 3E). These findings suggest that diminished dynamical range in between-network connectivity contributes to the observed cognitive impairments, and partially to performance recovery following a nap.

To probe the temporal coupling between brain connectivity and lapses in performance, we examined trial-level associations between the participation coefficient (BT) and lapse occurrences. A general linear model revealed that fluctuations in BT significantly tracked the occurrence of behavioural lapses across tasks in the well-rested condition (Figure 3F). The spatial distribution of regions showing positive and negative associations was aligned with the primary canonical gradient of functional connectivity ranging from transmodal (high BT) to sensory (low BT) regions (Margulies et al. 2016). Specifically, at the moment of cognitive lapses, BT was significantly high within the DMN. Importantly, this pattern was significantly diminished following Sleep Deprivation and the Recovery Nap, suggesting a reduced fluctuation of BT tracking cognitive lapses (Figure 3F).

Finally, we quantified the proportion of lapses occurring within integrated versus segregated modes. Sleep deprivation did not alter the likelihood that lapses occurred during integrated or segregated modes, and the likelihood of occurrences did not change following the nap (Figure 3G).

### Thalamic control of integration–segregation dynamics is disrupted after sleep deprivation

Thalamocortical (static) functional connectivity differed between conditions, with widespread reductions in correlations between the timeseries of the thalamus and 400 cortical parcels following sleep deprivation compared with rested wakefulness (t_(7999)_ = 69.2, p = 2.2 × 10^−16^), and recovery after the nap (t_(7999)_ = 63.8, p = 2.2 × 10^−16^ Figure 4A). Across participants, these reductions in static thalamocortical connectivity were positively correlated with changes in the variability of between-network connectivity (BTstd) in the DMN and Limbic networks, such that cortical regions showing greater reductions in BT variability also exhibited larger decreases in thalamic coupling (Figure 4B). Conversely, this relationship was opposite in the Dorsal Attention Network and Somatosensory networks (Figure 4B).

**Figure 4.**
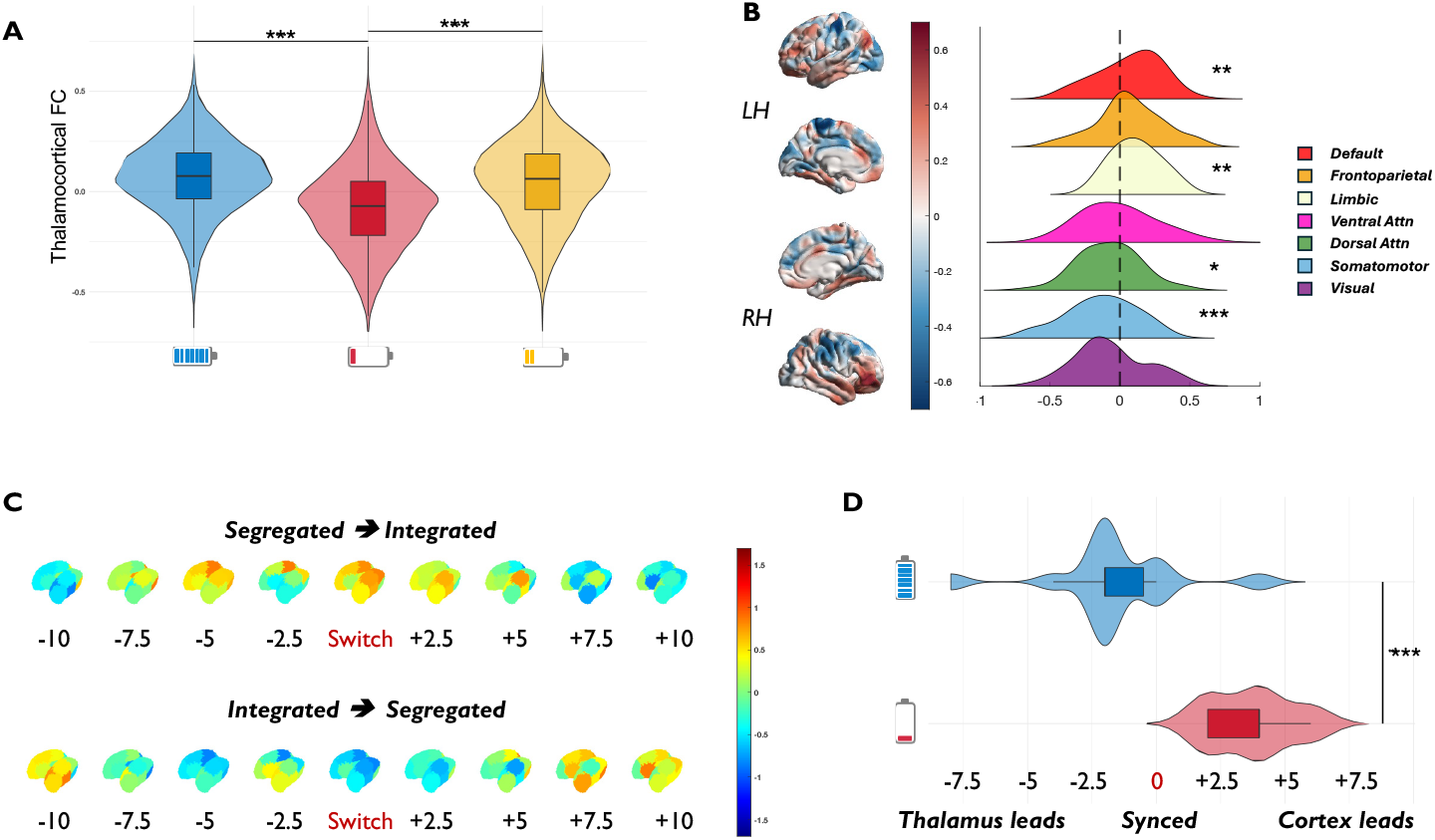
Thalamic control of dynamical integration modes is impaired following sleep deprivation. **A**| Thalamocortical functional connectivity during Rested Wakefulness, Sleep Deprivation, and Post-nap recovery. Plots show condition-wise mean differences in (static) functional connectivity between the (whole) thalamus and 400 cortical parcels. **B**| Parcel-wise correlation between SD-induced changes in BT temporal variability and static thalamocortical FC. The cortical map shows, for each parcel, the correlation across participants between ΔBTstd (SD–WR) and Δ thalamus–cortex static FC (SD–WR) (warm = positive, cool = negative, unthresholded map shown). Right, stacked kernel density plots summarise these correlation coefficients within each Yeo 7 network; dashed line marks r = 0. **C**| Lagged coupling between thalamic activity and transitions between network modes. A GLM related mode switches (integrated ↔ segregated) to thalamic time series across lags (−4 to +4 windows). Panels show group t-statistics mapped onto 14 thalamic nuclei for each lag during Rested Wakefulness (top: integrated→segregated; bottom: segregated→integrated); warmer colours indicate positive associations, cooler colours negative. **D**| Peak timing and strength of thalamic coupling to mode transitions. Sleep deprivation delayed the thalamic response to transitions between integrated and segregated network modes compared with Rested Wakefulness. Boxplots represent the temporal peak of coupling to state switches across all 14 thalamic nuclei.

To examine the temporal relationship between thalamic activity and large-scale integration– segregation dynamics, we modelled transitions between integrated and segregated modes as predictors of the time series from 14 thalamic subnuclei with general linear models. During rested wakefulness, activity of thalamic nuclei consistently peaked at or just before transitions from segregated to integrated modes, whereas activity across all nuclei was suppressed during switches from integrated to segregated modes (Figure 4C). After sleep deprivation, this coupling was weakened and shifted in time, indicating a delay in thalamic responses to network reconfigurations (Figure 4D). Together, these results suggest that sleep deprivation disrupts thalamic coordination of integration– segregation dynamics, reducing both the strength and temporal precision of thalamic control over cortical network states.

### Endogenous fluctuations are linked to altered integration dynamics after sleep deprivation

Parcel-wise analyses revealed regionally specific changes in the amplitude of BOLD signal fluctuations following sleep deprivation. Fluctuations decreased in default mode network (DMN) regions and increased in primary sensory areas, including visual and somatomotor networks (Figure 5A), consistent with the principal cortical gradient (Margulies et al. 2016). Across parcels, group-level fluctuation changes following sleep deprivation were negatively associated with changes in between-network connectivity, such that regions showing increased signal fluctuations were also those with greatest reductions in the temporal variability of the participation coefficient (ΔBTstd, ρ = –0.24, p < 0.001), an effect strongest within the DMN (Figure 5B).

**Figure 5.**
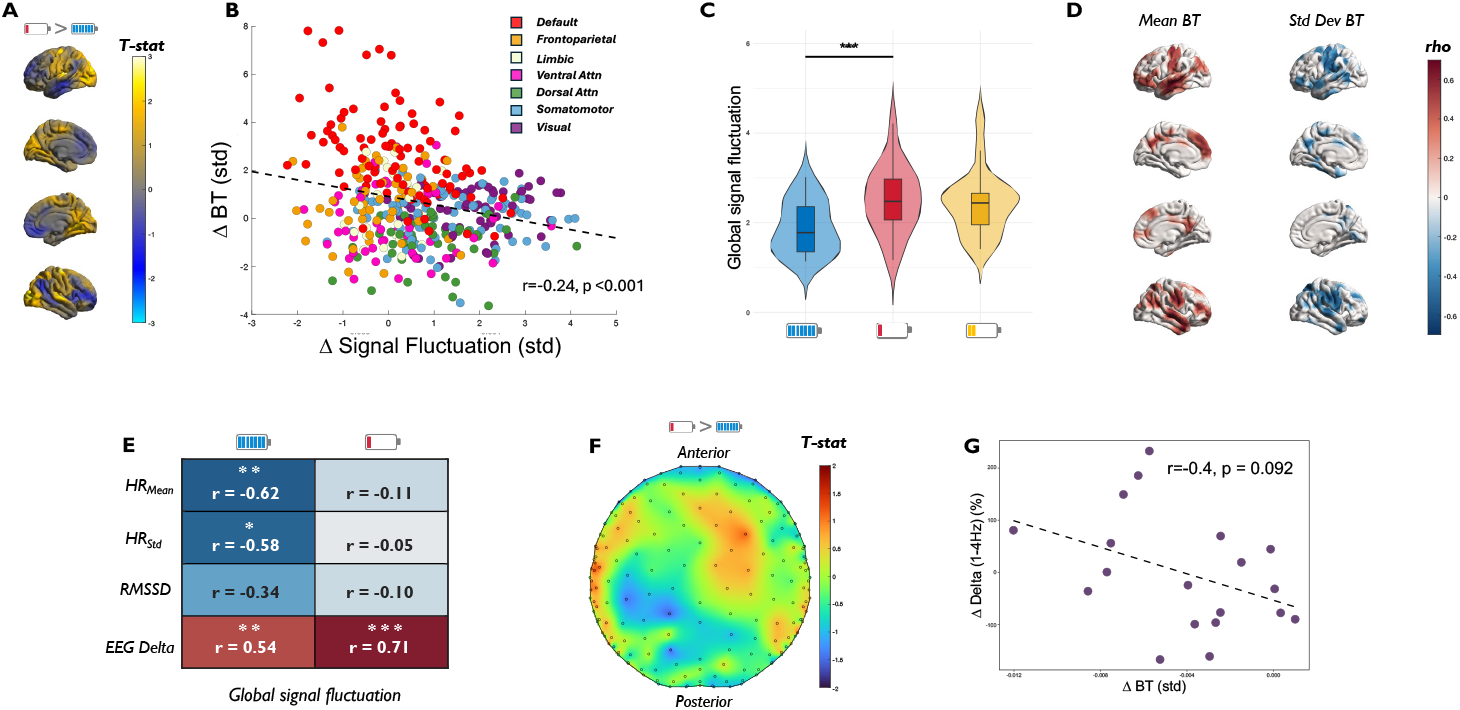
Relationship between endogenous fluctuations and configurations of integration brain connectivity. **A**| Regional amplitude fluctuations of cortical activity (parcel-wise BOLD standard deviation) comparing Rested Wakefulness and Sleep Deprivation. Surface maps show t-statistics for WR→SD differences across 400 parcels (warm = greater fluctuation after SD; cool = reduced). **B**| Relationship between regional amplitude fluctuations and between-network connectivity. Scatterplot shows parcel-wise associations between changes in BOLD signal variability (WR→SD) and changes in participation coefficient (BT). Each point represents a cortical parcel, coloured by Yeo 7 network; dashed line indicates the group-level regression fit. **C**| Global signal fluctuations across conditions. Violin plots show the amplitude of whole-brain BOLD signal variability during Rested Wakefulness, Sleep Deprivation, and Post-nap recovery. Black lines indicate the median across participants. **D**| Relationship between global signal fluctuations (std deviation of mean signal from gray matter) and between-network connectivity. Surface maps show parcel-wise correlations between WR→SD changes in global signal fluctuation and changes in participation coefficient. Left: mean BT (ΔBTave); Right: temporal variability of BT (ΔBTstd). **E**| Coupling between global signal fluctuations and physiological indices. Heatmap shows correlations between global BOLD signal fluctuation and ECG/EEG measures—mean heart rate, heart-rate variability (SD of HR), root mean of the successive differences of heart rate (RMSSD), and EEG delta power during Rested Wakefulness (left column) and Sleep Deprivation (right column). Warmer colours indicate stronger positive correlations; cooler colours indicate negative correlations. **F**| Scalp topography of WR→SD changes in EEG delta power (1–4 Hz). **G**| Relationship between change in EEG delta power and between-network connectivity variability (ΔBTstd).

At the whole-brain level, global signal fluctuations (standard deviation of the global signal time series from the entire gray matter, GSF) were elevated during sleep deprivation relative to rested wakefulness, with no significant differences in the post-nap condition (Figure 5C). Parcel-wise analyses showed that increases in GSF were positively related to increases in mean BT in somatomotor and DMN regions, and negatively related to changes in BT variability (Figure 5D).

Physiological associations with GSF were condition-dependent. In the well-rested state, GSF was strongly negatively correlated with mean heart rate (HR) and HR variability (SD of HR), and positively correlated with EEG delta power (1 – 4 Hz); the relationship with Root Mean Square of Successive Differences (RMSSD) was negative but did not reach significance. After sleep deprivation, these cardiac associations were abolished, while the positive correlation with EEG delta strengthened (Figure 5E). However, scalp-level EEG analyses revealed no significant change in delta power between conditions (Figure 5F), and inter-individual differences in delta power change were only weakly related to changes in BT variability (ρ = –0.40, p = 0.092; Figure 5G).

Together, these findings indicate that sleep deprivation alters both regional and global BOLD fluctuations, disrupts their normal coupling with cardiac physiology, and links elevated global synchrony to reduced flexibility of large-scale integration dynamics.

## Discussion

By combining time-resolved fMRI, EEG, and ECG measures during cognitive task performance, we demonstrate that sleep deprivation shifts network topology toward more intermediate connectivity states, characterised by increased between-network connectivity but reduced temporal variability (Figure 1). These alterations were expressed as a dampening of transitions between integrated and segregated brain modes (Figure 2), linked to impaired task accuracy and slowed responses (Figure 3). At the physiological level, sleep deprivation weakened thalamic coordination of network dynamics (Figure 4) and disrupted the normal coupling between cardiac and global brain fluctuations, while strengthening associations with sleep-like EEG delta activity (Figure 5). Together, these findings provide converging evidence that sleep deprivation compromises cognitive control through alterations in the brain’s capacity to flexibly reconfigure its large-scale functional architecture, driven by altered thalamocortical regulation and disrupted integration of central and peripheral physiological signals.

In the present study, sleep deprivation increased average participation coefficient (BT), indicating a more even distribution of connections between cortical networks, but did not alter the mean within-module degree (WT). The participation coefficient reflects the spread of a node’s connections across networks relative to its total degree and can rise even if absolute correlation strengths remain unchanged, provided that cross-network links become more prevalent. Such redistribution is consistent with recent work showing that fluctuations in arousal are accompanied by dynamic reconfiguration of hub topology, whereby high-participation “connector hubs” shift their network affiliations as vigilance declines (Lee et al., 2022). By contrast, the within-module degree z-score is normalised within each network and reflects a node’s relative standing in within-network connectivity; it remains stable if all nodes increase a similar proportion of within-network to cross-network edges. This pattern contrasts our earlier findings exploring static integration using information theory (Cross et al., 2021, PLOS Biology), where sleep deprivation increased global integration, but at a higher rate within canonical cortical networks than between them, resulting in functional clustering (or segregation). Integration in that context purely indexed the magnitude of statistical dependence, and did not capture network topology, only the strength of coupling within and between networks. The present results therefore suggest that, rather than just strengthening within-network coupling, sleep deprivation also redistributes connectivity toward more inter-network interactions without altering the relative within-network rankings of individual regions. This interpretation is consistent with our previous observation that large-scale functional organisation, as indexed by cortical connectivity gradients, remains stable after sleep deprivation (Cross et al., 2021, NeuroImage), suggesting that redistribution occurs within an overall preserved macroscale architecture.

Building on this distinction, our dynamic analyses also revealed that sleep deprivation not only redistributes connectivity across networks but also constrains the brain’s ability to flexibly alternate between integrated and segregated configurations. Whereas rested wakefulness was characterised by clear separation and stable occupancy of these two modes, sleep deprivation dampened the distinction between these modes and increased transitions between them, resulting in a contraction toward intermediate network states. This contrasts with findings of a reduction in the number of dynamic state transitions following sleep deprivation (Teng et al. 2019, NeuroImage). However, that approach identified five recurring connectivity states, whereas the present analysis focused specifically on two “extreme” modes of network topology (integrated versus segregated). The apparent discrepancy may therefore reflect methodological differences: rather than decreasing the number of overall state transitions, sleep deprivation may selectively compress the brain’s exploration of topological extremes. Thus, the global increase in integration observed at the static level appears to manifest dynamically as an instability of network topology, with reduced stability of both integrated and segregated modes and greater persistence in intermediate configurations.

The associations observed here between changes in the temporal variability of between-network connectivity (ΔBT_SD–WR_) and changes in behavioural performance closely parallel those we previously reported using static, information-theoretic measures of integration (Cross et al., 2021, *PLOS Biology*). This convergence suggests that while dynamic analyses capture the temporal unfolding of integration and segregation, both approaches are sensitive to the same underlying mechanism linking large-scale network integration to cognitive vulnerability under sleep loss. The present framework therefore extends our earlier findings by revealing that reduced behavioural performance after sleep deprivation is accompanied not only by higher static integration but also by a loss of temporal flexibility in network topology.

The present findings suggest that sleep deprivation disrupts the mechanisms that normally regulate the balance between integrated and segregated brain states. Under rested conditions, the thalamus appears to play a central role in this regulation: activity across thalamic nuclei reliably peaked just prior to transitions into integrated states and was suppressed during transitions into segregated states, consistent with a gating mechanism that helps to regulate cortical excitability through activity-dependent inhibitory mechanisms (Halassa & Sherman, 2019, Neuron). This aligns with the known role of intralaminar and midline thalamic nuclei, which receive dense input from brainstem arousal centres and project diffusely to the cortex (Edlow et al., 2012, J Neuropathol Exp Neurol; Muller et al., 2020, NeuroImage), supporting arousal (Chang et al. 2016, PNAS) and coordinating widespread functional modes. After sleep deprivation, this regulatory influence was both weakened and delayed, pointing to impaired thalamocortical gating of network transitions.

In parallel, cortical fluctuations became more pronounced in sensory networks and in the global signal, an instability in vigilance and arousal regulation that has been observed in states of drowsiness (Wong et al., 2013, NeuroImage) and light sleep (Larson-Prior et al., 2009, PNAS). In the well-rested state, these global fluctuations are coupled to cardiac physiology (mean HR and HR variability), but after sleep deprivation this relationship was lost, suggesting that ascending autonomic influences on the cortex may become uncoupled. At the same time, global fluctuations became more strongly tied to EEG delta activity, consistent with a shift toward sleep-like dynamics. Previous EEG–fMRI studies similarly report increased global signal amplitude during low vigilance states (Wong et al., 2013, NeuroImage) and positive delta-DMN connectivity during sleep inertia (Vallat et al., 2018, NeuroImage), while broadband EEG power fluctuations generally track global fMRI dynamics across arousal states (Wen & Liu, 2016, J Neurosci). The patterns observed here thus appear to reflect a transitional, unstable regime of reduced arousal, more akin to light sleep or drowsiness than to consolidated sleep (Horovitz et al., 2008; Tagliazucchi & Laufs, 2014). Taken together, these observations suggest that sleep deprivation constrains the brain’s ability to flexibly alternate between modes of integration and segregation by simultaneously weakening thalamic regulation and altering the coupling between cortical and peripheral physiological signals.

Several limitations should be acknowledged when interpreting these findings. First, the sample size was modest (n = 20), and replication in larger cohorts will be important to establish the robustness and generalisability of the effects. Nonetheless, the bidirectional design of the study enhances the reliability of the findings. Second, the simultaneous acquisition of task-based fMRI with EEG and ECG provided valuable multimodal insights but also introduces complexity. It remains difficult to disentangle task-specific demands from broader fluctuations in vigilance and arousal, and future work using resting-state paradigms or multimodal replication could help clarify this distinction. However, resting state fMRI studies of sleep deprivation are challenging given the heightened propensity to fall asleep under resting conditions. Third, our analytical approach relied on a sliding-window method and k-means clustering into two modes; while this approach is well-established, alternative techniques such as hidden Markov models (Quinn 2018), co-activation pattern analysis (Karahanoğlu 2015), or point-process methods (Wu 2015) may provide complementary perspectives on the temporal organisation of network dynamics. Given the non linear relationship between brain activity and hemodynamic fluctuations, interpreting temporal associations between behaviour and physiological responses should be made with caution. Fourth, although we included ECG measures, we did not directly record respiration, which is known to contribute to global BOLD fluctuations. Estimating respiratory parameters from ECG or incorporating direct respiratory measures could refine interpretations of the observed global signal effects. Similarly, we did not model cardiac and respiratory signals explicitly (e.g., RETROICOR/RVT); although our preprocessing pipeline mitigated physiological contamination, residual respiratory variance may have persisted and could have contributed to condition differences in global (Yuan et al. 2025) or regional (Yuan et al. 2016) signal fluctuations. Finally, future studies should move beyond characterisation to probe causal mechanisms, for example, by manipulating arousal pharmacologically (Gordji-Nejad et al. 2024, Scientific Reports) or via targeted sleep interventions (Perrault et al. 2025, medRxiv), extending designs to longer periods of sustained wakefulness, or examining the generalisability of these effects to clinical and at-risk populations such as shift workers, individuals with hypersomnolence, or ageing cohorts.

This study advances our understanding of how sleep deprivation alters brain function by demonstrating that the loss of cognitive control is accompanied by a narrowing of the brain’s dynamical repertoire, linking network topology, thalamic regulation, and global brain–body fluctuations. Together, these findings establish disrupted flexibility of integration and segregation as a key mechanism of vulnerability to sleep loss, highlighting dynamic connectivity as a sensitive marker and potential target for mitigating cognitive decline in real-world and clinical contexts.

## Methods

### Population and Experimental Design

Twenty participants were recruited using advertisements posted online and within Concordia University, Montreal. This study was approved by the Comité central d’éthique de la recherche (CCER), established by the Ministère de la Santé et des Services sociaux in Quebec. This ethical review of research adheres to the principles expressed in the Declaration of Helsinki. Informed written consent was obtained from all participants. All participants were 18-30 years old, healthy, regular sleepers and none were taking any medication. Participants made three visits to the sleep laboratory, including a night for habituation and screening to rule out sleep disorders, a normal night (8hr sleep, RW) and experimental night (0hr sleep, SD). Each morning in the RW and SD conditions, participants completed one resting state sequence and three cognitive tasks: the Attention Network Task, Mackworth Clock Task, and the N-back task. In the SD condition, participants were provided a 60-min opportunity to sleep inside the MRI (recovery nap), and the tasks and then the resting state sequence were repeated post nap, PN). The order of the RW and SD sessions were counterbalanced across subjects. The protocol has been described in more detail elsewhere (Cross et al., 2021, NeuroImage).

### Cognitive performance

All tasks were run on a laptop computer using Inquisit software (Millisecond Software LLC, 1998), displayed to the participant via a projector screen behind the MRI scanner. The participant responded to all tasks via button presses made using a response pad attached to the fingers of the left hand. For all tasks, outcomes of reaction time (ms) and accuracy (%; correct trials/number of trials) were measured. For the context of this study, lapses were defined as trials consisting of missed or incorrect responses.

For the change in performance outcomes between conditions (WR, SD, PN), each of the 3 separate cognitive tasks were grouped into either accuracy (% of correct responses) or speed (reaction time, in ms). This provided 2 data matrices of size #subjects x #tasks; one matrix containing accuracy scores and one containing reaction times. The change scores for accuracy or speed were entered into separate Principal Component Analyses (PCA), in order to provide a global estimate of performance. The first principal component from each PCA was extracted and used as a single measure for Accuracy or Speed across the 3 tasks (as per Cross et al 2021b).

### fMRI Data Acquisition and Analysis

EEG and fMRI data acquisition and preprocessing have been described in detail in our previous studies (Cross 2021a, Cross 2021b, Uji et al 2022). MRI scanning was acquired with a 3T GE scanner (General Electric Medical Systems, Wisconsin, US) using an 8-channel head coil. Functional scans were all acquired using a gradient-echo echo-planar imaging (EPI) sequence (TR = 2500 ms, TE = 26 ms, FA = 90°, 41 transverse slices, 4-mm slice thickness with a 0% inter-slice gap, FOV = 192 × 192 mm, voxel size = 4 × 4 × 4mm3 and matrix size = 64 × 64). Structural T1-weighted images with a 3D BRAVO sequence, and functional echo planar images were acquired. Data were preprocessed using the fMRIPrep (Esteban et al., 2019) and XCP-D (Ciric et al., 2017) toolboxes (for detailed steps, see Cross et al. 2021, PLOS Biology). The preprocessed BOLD time series for each subject was projected onto the cortical surface and smoothed along the surface using a 6mm smoothing kernel using the Freesurfer software package.

To control for spurious patterns of connectivity associated with task-evoked activity, the general task-specific activity (onset stimuli convolved with a hemodynamic response function) was first regressed out from the BOLD time series, similar to (Shine et al. 2016). The time series from all tasks were then concatenated into one timeseries per subject and condition to. This resulted in a total timeseries of 26 minutes (624 TRs) per condition (WR, SD & PN).

### Network and assembly identification

For all analyses of task data, the BOLD time series for each vertex on the fsaverage surface was assigned to one of 400 cortical parcels from a pre-determined standardised template of functionally similar cortical regions (Shcaeffer 2018). The time series for all the vertices corresponding to each parcel were averaged to give one mean time series per parcel, resulting in a total of 400 time series across both cortical hemispheres. We also extracted timeseries from 7 regions of the thalamus, using the HCP thalamic atlas (Najdenovska 2018) as provided by XCP-D.

Time-resolved functional connectivity was calculated between all 400 cortical regions using the multiplication of temporal derivatives (MTD) metric (Shine et al. 2015). The MTD is computed by calculating the point-wise product of temporal derivative of pairwise time series, then averaged over a temporal window to reduce the contamination of high-frequency noise in the time-resolved connectivity. The MTD was calculated within a sliding temporal window of 10 time points (∼25s) with a step of 2.5s, balancing temporal resolution against the reliability of connectivity estimates (Shine et al. 2015). Individual functional connectivity matrices were then calculated within each temporal window.

### Time-Resolved Network Topology and Community Structure

To examine dynamic changes in brain network organisation, we estimated both time-averaged and time-resolved community structure using the Brain Connectivity Toolbox (Rubinov & Sporns, 2010). For each cortical parcel, we computed two key graph-theoretical metrics within each temporal window: the module-degree z-score (within connectivity, WT) and the participation coefficient (between connectivity, BT). The WT quantifies rank-based within-module connectivity, indicating how strongly a node is connected to other nodes in its own module. The WT Z-scores were calculated as the number of connections each node made to other nodes within the same module, normalized (z-scored) relative to the mean and standard deviation of within-module degrees across all nodes in that module (Guimerà & Amaral, 2005). A positive within-module degree Z-score indicates that a node is more connected within its own module than average (a local hub), whereas a negative Z-score indicates fewer within-module connections than most nodes in that module. The BT measures between-module connectivity, reflecting how evenly a node distributes its connections across different modules. A BT value close to 1 suggests a region has broad, integrative connectivity across modules, whereas a value near 0 indicates that its connections are largely confined within a single module. Modules were set as the Yeo 7 networks (Yeo et. al 2007) and were fixed for every time window. Graph metrics were computed on weighted, undirected matrices.

To characterise fluctuations in global network topology over time, firstly we assessed variability by calculating the standard deviation of BT and WT over time for each participant and session. We additionally used the method from (Shine et al., 2016) and constructed a “Cartographic Profile” for each condition. For each time window, we created a two-dimensional joint histogram of WT and BT scores across all regions (101 × 101 unique bins, equally spaced between 0 and 1 for BT and –5 and 5 for WT), capturing the distribution of the 400 parcels in terms of their within- and between-module connectivity. Each bin of this histogram represents a specific functional configuration (WT:BT) that a cortical parcel could exhibit in the given time-window. Profiles skewed toward lower BT and higher WT values (top left on the 2D joint histogram) reflect segregated network configurations, whereas higher BT and WT values (top right on the joint histogram) correspond to integrated configurations. These cartographic profiles were used to track time-varying shifts in the brain’s topological state throughout the scanning period. By concatenating the 2D joint histograms across all time-windows, it is possible to see which configurations are traversed most frequently across time, with colour intensity indicating the number of parcels occupying that configuration across time.

For each participant (n=20) and experimental session (rested wakefulness, sleep deprivation, and recovery nap), we extracted time-resolved values of the participation coefficient (BT) or within-module degree (WT) from the pre-computed graph metrics (400 parcels). At the first level, paired t-tests were performed within each subject for each parcel across all temporal windows, comparing (i) sleep deprivation versus rested wakefulness, and (ii) recovery nap versus sleep deprivation. This yielded a parcel-wise distribution of t-statistics per subject for each contrast of interest. At the second level, we tested whether these subject-specific effects were consistent across the group. For each parcel, the distribution of first-level t-statistics across all subjects was entered into a one-sample t-test, separately for the sleep deprivation > rested wakefulness and recovery nap > sleep deprivation contrasts. Group-level t-values and corresponding p-values were retained for each parcel, producing a parcel-wise statistical map of condition effects on BT or WT. Multiple comparisons were controlled across the 400 parcels using a false-discovery rate (FDR) method (Benjamini and Hochberg, 1995).

To compare how the distribution of time-resolved network configurations differed across conditions (WR, SD, PN), we quantified the proportion of time windows that each subject spent within every bin of the two-dimensional cartographic profile (WT × BT space). These proportions were normalised within each subject to sum to 1, yielding a probability distribution of occupancy across the cartographic space. A mixed-effects general linear model was then used to test for bins showing significant modulation by condition, accounting for repeated measures across participants. Pairwise contrasts between conditions were assessed with paired-sample t-tests on the occupancy proportions for each bin, and statistical significance was determined using a false discovery rate (FDR) correction of q < 0.05.

Finally, to test whether the cartographic profile of the resting brain fluctuated over time between two topological extremes (ie. global integration vs segregation), we performed clustering of temporal windows. Specifically, we classified the joint histogram of each temporal window (which is naive to cartographic boundaries) over time using a k-means clustering analysis (k = 2). As a result of this analysis, each window was assigned to one of two clusters – corresponding to the extremes of integrated and segregated brain modes (see Supplementary Figure 1.2). Clustering was performed specifically for each subject and condition.

### Linking between-network connectivity to behavioural lapses

To test whether moment-to-moment changes in between-network connectivity tracked behavioural failures, we examined the relationship between the participation coefficient (BT) and the occurrence of task lapses. Behavioural lapses were convolved with a 10-TR sliding window to align with the temporal resolution of the dynamic connectivity estimates. A general linear model (GLM) was then fitted to relate lapse time series to parcel-wise BT fluctuations centered around lapse onset. This allowed estimation of how strongly parcel connectivity dynamics were associated with behavioural lapses. Group-level effects were assessed by performing one-sample t-tests on subject-level regression coefficients. Spatial maps of significant associations were generated by projecting parcel-level t-statistics onto the cortical surface.

### Thalamic tracking of integration–segregation dynamics

To assess the temporal relationship between thalamic activity and large-scale network reconfigurations, we constructed a regressor marking transitions between integrated and segregated modes (derived from k-means clustering of cartographic profiles, k = 2). General linear models (GLMs) were used to predict thalamic and cortical time series from the mode-switch regressor across temporal lags (−4 to +4 windows, sliding by one TR). For each lag, regression coefficients were estimated per subject and region, and group-level one-sample t-tests were performed across participants. Thalamic nuclei were defined using the HCP 14-region atlas, and cortical parcels used the 400-region Schaefer parcellation. Lag profiles for each nucleus were smoothed with a 5-TR (12.5 sec) moving average to aid interpretability. Peak lag and effect size were compared between WR and SD to quantify shifts in thalamic engagement with network transitions.

### Physiological Data Acquisition and Analysis

EEG was acquired using an MR compatible 256 high-density geodesic sensor EEG array (Electrical Geodesics Inc, Magstim EGI). EEG data were recorded at 1000Hz referenced to Cz using a battery-powered MR-compatible 256-channel amplifier. Electrocardiography (ECG) was also collected via two MR compatible electrodes through a bipolar amplifier (Physiobox, EGI). ECG was visually inspected and R-peaks were detected in a semi-automatic manner (automated detection manually reviewed and corrected). The EEG data were then corrected for MR gradient and ballistocardiographic pulse-related artefacts using the Brainvision Analyzer (Brain Products Inc, Gilching Germany) as per previous papers (Uji et al. 2022). The MR-denoised EEG signal was bandpass filtered between 1 and 20 Hz to remove low-frequency drift and high-frequency noise, down-sampled to 250 Hz, and re-referenced to the linked mastoids. For the analysis of wake EEG, eye blink artefacts were removed with ICA using the MNE Python package (Gramfort et al. 2014). The ICA was fit on each EEG timeseries concatenated across tasks for each subject and condition. The number of components was set to 15, with a random seed to ensure the uniformity of components. The resulting ICA components were then compared to EOG electrodes any ICs that matched the EOG pattern were automatically marked for exclusion. Any matching components were then excluded from the original signal, and this new timeseries was then used for all subsequent analysis. Power spectral analysis was also computed using MNE, via the Welch method using a frequency resolution of 1Hz. Analysis of heart rate from ECG were performed using the ‘sleepecg’ Python package. Measures of mean HR, HR variability (standard deviation of HR) and Root Mean Square of Successive Differences (RMSSD) were extracted from the ECG.

---

**Supplementary Figure 1.**
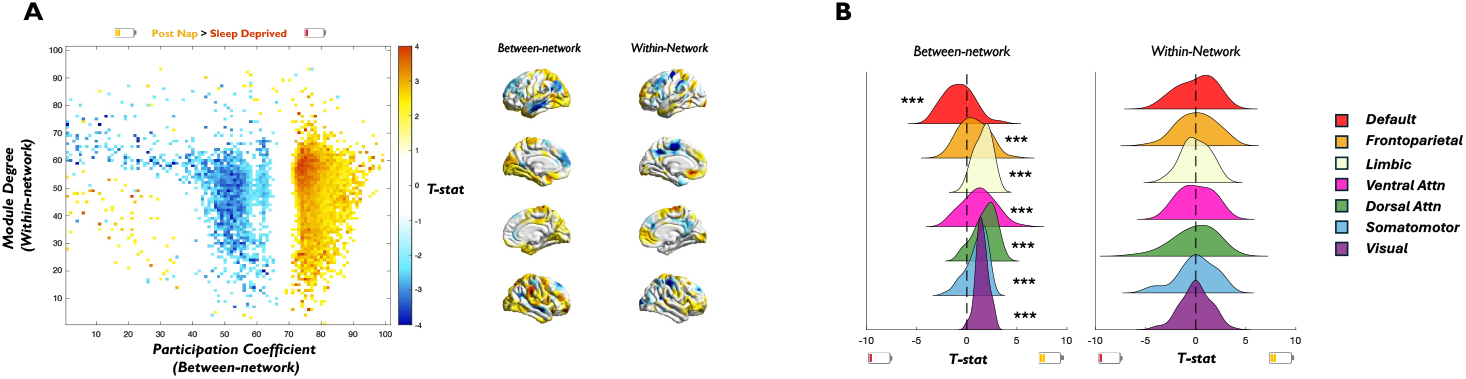
Changes in configurations of integration and segregation after a recovery nap. **A** | Surface projections of cortical parcels associated with higher Between-network (left) or Within-network (right) connectivity during the Post-Nap state, when compared the Sleep-Deprived state. **B** | Changes in Between-network connectivity within the 7-Yeo functional networks demonstrated a significant decrease in DMN, with significant increases in all other networks.

**Supplementary Figure 3.**
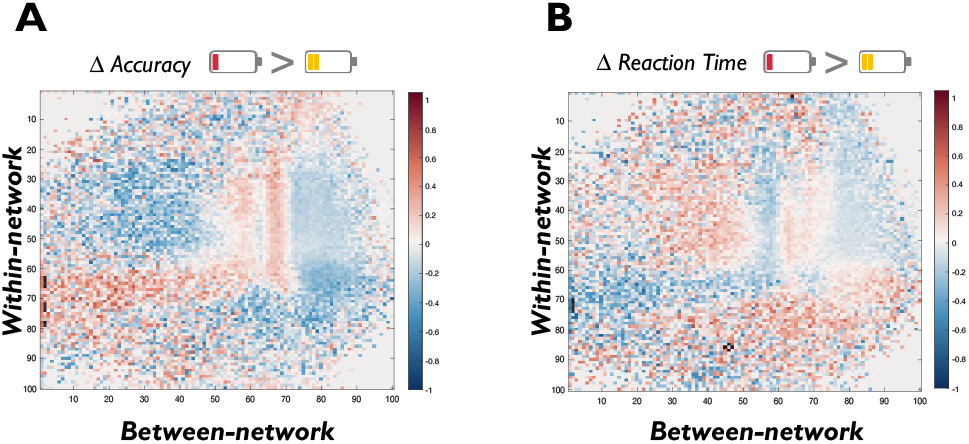
Association between reconfigurations of cartographic profile and cognitive performance following a recovery nap. **A**| Differences in the cartographic profile between Sleep Deprivation and Post Nap correlated with changes in task accuracy. The heatmap shows correlation coefficients between topological reorganisation (WT–BT joint space) and behavioural scores, with warm colours indicating positive associations and cool colours negative associations. Outlined regions indicate clusters surviving significance thresholding (p < 0.05, FDR corrected). **D**| Correlation between reconfiguration of brain network dynamics and changes in task reaction time, displayed as in (A).

